# Deleterious mutations can contribute to the evolution of recombination suppression between sex chromosomes

**DOI:** 10.1101/2024.11.18.624013

**Authors:** Paul Jay, Amandine Véber, Tatiana Giraud

## Abstract

Many organisms have sex chromosomes with large non-recombining regions that expand in a stepwise manner, although the underlying reasons remain poorly understood. Recently, we proposed that recombination suppression may evolve in sex chromosomes simply because of the presence of recessive deleterious mutations within genomes. Specifically, we demonstrated that chromosomal inversions suppressing recombination and carrying by chance fewer deleterious mutations than average have a selective advantage. In addition, we showed that the permanent heterozygosity of Y-like sex chromosomes facilitates the fixation of these less-loaded inversions by a sheltering effect, i.e., by preventing the expression of recessive deleterious mutations in a homozygous state when they increase in frequency. In contrast, similar less-loaded inversions in autosomes suffer from a disadvantage in the homozygous state as their frequency increases, preventing their fixation. However, the methodology and significance of our previous study have been questioned. Here, we present new analyses that further reinforce our original claims, demonstrating that the lower-load advantage and the sheltering effect can explain the fixation of inversions on sex chromosomes over a broad range of parameter values. We show that these mechanisms promote the fixation of inversions on Y-like chromosomes at rates exceeding those expected under drift alone. We used, as a control, autosomes with a similar population size as the Y chromosome, which, we argue, provides the appropriate neutral control for the sheltering effect. We also address criticisms regarding our focus on inversions surviving the first 20 generations in a figure of our previous study, and show that these criticisms stemmed from a misunderstanding of what this figure was intended to illustrate. Including all inversions, even those that went extinct within 20 generations, does not alter our conclusions. Overall, the present study offers new support for our theory based on the combination of lower-load advantage and sheltering effect, and addresses the questions about its significance and range of applicability.

---

Many organisms have sex chromosomes with large non-recombining regions that expand in a stepwise manner, although the underlying reasons remain poorly understood (Ponnikas et al., 2021, Wright et al. 2016, Jay et al. 2024; Saunders and Muyle 2024). It has long been considered that recombination suppression on sex chromosomes gradually expands because selection favors the linkage of sexually antagonistic loci to sex-determining genes (Ruzicka & Connallon, 2020, Wright 2016, Rice 1987). However, no studies have been able to demonstrate that this mechanism is really responsible for evolutionary strata in natural populations so far (Ponnikas et al., 2021, Ironside 2010, Beukeboom and Perrin, 2004, but see Wright et al., 2017) and recombination suppression has also been reported to gradually expand around many fungal mating-type loci and other supergenes despite the lack of sexual antagonism (Hartmann at al. 2021, Branco et al., 2017, Wang et al., 2013, Jay et al., 2024, 2021, Yan et al., 2020). In addition, theoretical issues have been raised about the model of sexual antagonism driving the evolution of sex chromosomes (Cavoto et al., 2018). Altogether, this suggests that other mechanisms can drive the stepwise extension of recombination suppression (Jay et al., 2024).

Recently, we developed a general model for testing the idea that recombination suppression may extend stepwise around sex-determining or mating-type loci simply because of the presence of partially recessive deleterious mutations in genomes (Jay et al., 2022). In brief, the stochastic nature of deleterious mutation arrangement generates variations in fitness in natural populations, which can lead to the rise in frequency of non-recombining fragments (e.g. chromosomal inversions) carrying fewer deleterious mutations than average in the genomic region, i.e. being *less loaded*. However, if a less-loaded inversion is located on an autosome and the deleterious mutations are at least partly recessive, its selective advantage should decrease and eventually disappear as it increases in frequency in the population. Indeed, when the inversion becomes frequent enough to occur at the homozygous state, it expresses its recessive mutational load. In contrast, when such a less-loaded inversion is, by chance, fully linked to a permanently heterozygous allele (e.g. if it captures a Y-like sex-determining gene), its recessive deleterious mutations are *sheltered*, allowing the fixation of the inversion in the population of Y-like chromosomes, and thereby the formation of a genomic region lacking recombination between X-like and Y-like sex chromosomes. These mechanisms are thoroughly described and illustrated in Jay et al. (2024).

The successive fixation events of additional inversions linked to this first Y-linked inversion, by the same process, should lead to the further expansion of the non-recombining region between sex chromosomes, thereby forming multiple evolutionary strata of distinct levels of differentiation. It is important to note that the lower-load advantage and the sheltering effect are two distinct phenomena. The lower load is a fitness advantage, driving the increase in frequency of inversions and other types of non-recombining fragments. The sheltering effect only offsets the disadvantage of becoming frequent for an inversion carrying (partly) recessive deleterious mutations, allowing its fixation despite its load. In our previous study (Jay et al. 2022), we have proposed that the lower-load advantage and the sheltering effect can explain the evolution of sex chromosomes by their combined effects, but it is important to note that the sheltering effect could also play a role combined to other types of intrinsic advantages of inversions. Numerous other types of selective advantage can indeed be associated to inversions (e.g. sexual antagonism, local adaptation or beneficial effects of the inversion breakpoint, Jay et al. 2024). Regardless of the nature of their intrinsic advantage, inversions will suffer from homozygous disadvantage when increasing in frequency if they carry recessive deleterious mutations, unless they are Y-linked (Jay et al., 2022, 2024).

Following its publication (Jay et al., 2022), our theory generated significant interest but also faced criticisms (Olito and Charlesworth 2023, Charlesworth and Olito 2024). In the preprint entitled “Do deleterious mutations promote the evolution of recombination suppression between X and Y chromosomes?” (Olito and Charlesworth 2023), whose content was later published in a larger review on recent models of the sheltering hypothesis (Charlesworth and Olito 2024), the authors questioned the methodology and significance of our study. The criticisms focused on two methodological procedures: i) the removal of the simulations in which the inversions do not survive the first 20 generations in some figures, and ii) the lack of comparison with simulations with only neutral mutations (with all selective coefficients being zero, *s*=0).

Here, we present new analyses showing that these criticisms are unfounded and that our conclusions fully hold. The concern regarding the removal of the simulations in which the inversions do not survive the first 20 generations in the Figure 3C of Jay at al. (2022) stems from a misunderstanding of the objective of this Figure 3C. It is actually a direct consequence of the view of Olito and Charlesworth (2023) that our simulations should be compared to a case without any deleterious mutations, while the figure was intended for another comparison. Indeed, the Figure 3C in Jay et al. (2022) aimed at illustrating the comparison of inversion fixation probabilities between Y chromosomes and autosomes when the sheltering mechanism acts, i.e. when inversions become frequent (which we translated into “after 20 generations”). For this comparison and to illustrate solely the sheltering effect, the way of plotting this figure was fully valid and was explained in the legend.

Below, by re-analyzing Jay et al. (2022) dataset and performing new individual-based simulations, we show that that the removal of the inversions not surviving the first 20 generations does not change anything to our conclusions: the sheltering effect can protect inversions on the Y chromosome from expressing their recessive load, and can thereby, in combination with the lower-load advantage, lead to the progressive cessation of recombination on sex chromosomes, for a wide range of parameter values. In contrast, Olito and Charlesworth (2023, 2024) utilized our Figure 3C from Jay et al. (2022) to compare, on one hand, the rate of fixation of loaded inversions conditional on survival through the first 20 generations, and, on the other hand, the fixation probability of fully neutral inversions without any segregating deleterious mutations from generation 1. This comparison indeed makes no sense but the Figure 3C from Jay at al. (2022) was not intended for such a comparison, so that the innuendo of being misleading was unfair. The early dynamics of inversions (the first 20 generations) was studied in other figures in Jay at al. (2022).

The criticism regarding the lack of comparison with simulations without any deleterious mutations anywhere in genomes arises from contrasting views on what constitutes an appropriate “neutral control” for studying the impact of deleterious mutations on inversion fixation, which is an interesting and non-intuitive debate. Neutral controls are typically used in evolutionary biology to determine if a given advantage has a stronger effect than drift alone. We argue here that the appropriate control for drift, when considering the sheltering effect or the lower load advantage, is to examine inversions on autosomes with an effective population size equivalent to that of the Y chromosome, rather than a scenario devoid of deleterious mutations. Indeed, excluding deleterious mutations not only removes the sheltering effect but also removes the deleterious mutations that the sheltering effect mitigates, making it impossible to assess the impact and significance of the sheltering effect. Similarly, the lower-load advantage results from a change in the fitness landscape due to the presence of deleterious mutations. Thus, the question is not whether inversion fixation is more frequent when deleterious mutations segregate versus when they do not, but rather whether deleterious mutations, when present, significantly contribute to the fixation of new inversions, i.e. have a stronger effect in promoting inversion fixation than drift alone, all else being equal (i.e. with also the deleterious effect of mutations acting and preventing some inversion fixation). Below, we show, by comparing inversion fixation rates in autosomes and in sex chromosomes under various population sizes, that the sheltering effect significantly contributes to the increased fixation rate of inversions on Y sex chromosomes, i.e. that the higher fixation rates of inversions on the Y chromosome is not only due to its reduced effective population size.

## Material and methods

In this paper, we thus aimed at comparing inversion fixation probabilities between Y chromosomes and autosomes, including cases with similar effective population sizes between the two types of chromosomes, as a proper control for genetic drift, i.e., everything else being equal. We also further investigated the role of the lower-load advantage in driving the increase in frequency of inversions. To achieve these objectives, we used two datasets: the original simulation dataset from Jay et al. (2022) and a new set of individual-based simulations designed to extend this dataset with new parameter values, in particular allowing to compare sex chromosomes and autosomes with the same effective population size. We describe below in detail only the methods used to perform the new simulations. Each figure legend specifies the dataset used.

### New simulations

We used SLiM V4.1 (*80*) to simulate the evolution of panmictic populations of *N*=3,125 or *N*=12,500 individuals under a Wright-Fisher model. To assess the fate of inversions under various conditions, we simulated diploid individuals with a pair of 5Mb chromosomes representing, depending on the simulation, either autosomes or sex chromosomes. For the simulation of sex chromosomes, a single segregating locus with two alleles at the chromosome center was subject to balancing selection: one of the two alleles was permanently heterozygous, mimicking a classical XY system. Mutations occurred on chromosome at a rate *u=*1.2*10^−8^ per bp per generation and recombination occurred at a rate r=1.2*10^−8^ per bp per generation. Each new mutation had its selection coefficient *s* drawn from a gamma distribution with a shape of 0.2 and a mean of -0.01 or -0.001. The dominance coefficient of each mutation (*h)* was randomly sampled with uniform probabilities among either [0.1, 0.2, 0.3, 0.4, 0.5] (mean=0.3, no fully recessive mutations), or [0.0, 0.01, 0.1, 0.2, 0.3, 0.4, 0.5] (mean=0.22, includes full recessive mutations).

For each parameter combination (*h, s, N*, chromosome type), a simulation was run for 200,000 generations, to allow the population to reach an equilibrium for the number of segregating mutations. At the end of this initialization phase, the nucleotide diversity of populations ranged from π=0.000115 (with *s*=-0.01 and no fully recessive mutations) to π=0.000312 (with s=-0.001 and fully recessive mutations). Considering only the deleterious mutations with effects stronger than drift (i.e, with *s*<-1/N), diploid individuals carried on average one deleterious mutation every 10,000-15,000 base pairs. These levels of deleterious mutation densities are well within the order of magnitude of those estimated in natural populations: for instance, in humans, there is one heterozygous site every 1000bp, out of which about 25% have been estimated to be deleterious (Racimo and Schraiber, 2014). The population state was saved at the end of the initialization phase.

These saved states (one for each parameter combination) were repeatedly used as initial states for studying the dynamics of chromosomal inversions. Recombination modifiers mimicking inversions of 2Mb and 5Mb were then introduced at the center of the autosome or of the Y chromosome. These inversions suppress recombination across the region in which they reside when heterozygous. We first simulated inversions that were fully neutral in themselves (s_Inv_=0.0), their fitness only depending on the mutations they capture. In order to make the analysis of the sheltering effect more general, we then also simulated inversions associated with a fitness advantage (s_Inv_=0.01), that could for example be generated by the inversion breakpoint. For each parameter combination (*h, s, N*, chromosome type, chromosomal inversion size, inversion fitness), we ran 100,000 independent simulations, starting with the introduction of a single inversion in a single individual randomly sampled in the saved initial population. We monitored the frequency of these inversion-mimicking mutations during 25,000 generations, during which all evolutionary processes (such as mutation and recombination) remained unchanged, e.g. mutations were still appearing on inversions following their formation. Note that, when it occurs, an inversion captures deleterious mutations that are segregating in the population; such mutations could therefore also be present in several individuals without inversion in the populations. Our simulations thus do take into account the possibility of the presence of the same deleterious mutations on proto-X and proto-Y chromosomes during inversion spread, so that deleterious mutations on inversions on the Y chromosome are expressed in a homozygous state a rate equal to their frequency on X chromosomes. In order to reduce simulation time, simulations were stopped when inversions reached fixation, i.e. when the inversion reached a frequency of 1.0 on autosomes or 0.25 on sex chromosomes (i.e. 1.0 on Y-like chromosomes). Simulations were parallelized with GNU Parallel and plot were made with ggplot2. All scripts are available at https://github.com/PaulYannJay/MutationShelteringTheory/tree/main/Reanalyses_bioRxiv.

### Simulations from Jay et al. (2022)

The main differences in Jay et al. (2022), compared to the new simulations described above, were as follows: simulations were performed where mutations occurring on the chromosomes had all the same selection coefficients (s = −0.001, −0.01, −0.1, −0.25, or −0.5) and dominance coefficients (h = 0, 0.01, 0.1, 0.2, 0.3, 0.4, or 0.5). Additionally, simulations were conducted with a population size of N = 1,000 individuals, and the mutation rate was either u = 1e-08 or u = 1e-09.

## Results

Reanalysis of our simulations in finite populations with deleterious mutations segregating performed in Jay et al. (2022) showed that, across most of the parameter space examined, inversions were significantly more likely to spread and fix when they captured the sex-determining allele on the Y chromosome than when located on autosomes (Figure 1A). Inversion fixation rates on sex chromosomes exceeded those on autosomes across 94.9% of the parameter space explored, with inversions on Y chromosomes being, on average, 5.81 times more likely to fix than on autosomes across the whole range of parameter values studied. Considering only scenarios with mutations of realistic selection coefficients (e.g., s=[0.01,0.001]) and relatively recessive effects (e.g., h=[0,0.01,0.1,0.2]), inversions were 9.59 times more likely to fix on Y chromosomes than on autosomes. Additionally, inversions fixed 3.90 times faster on Y chromosomes than on autosomes.

**Figure 1.**
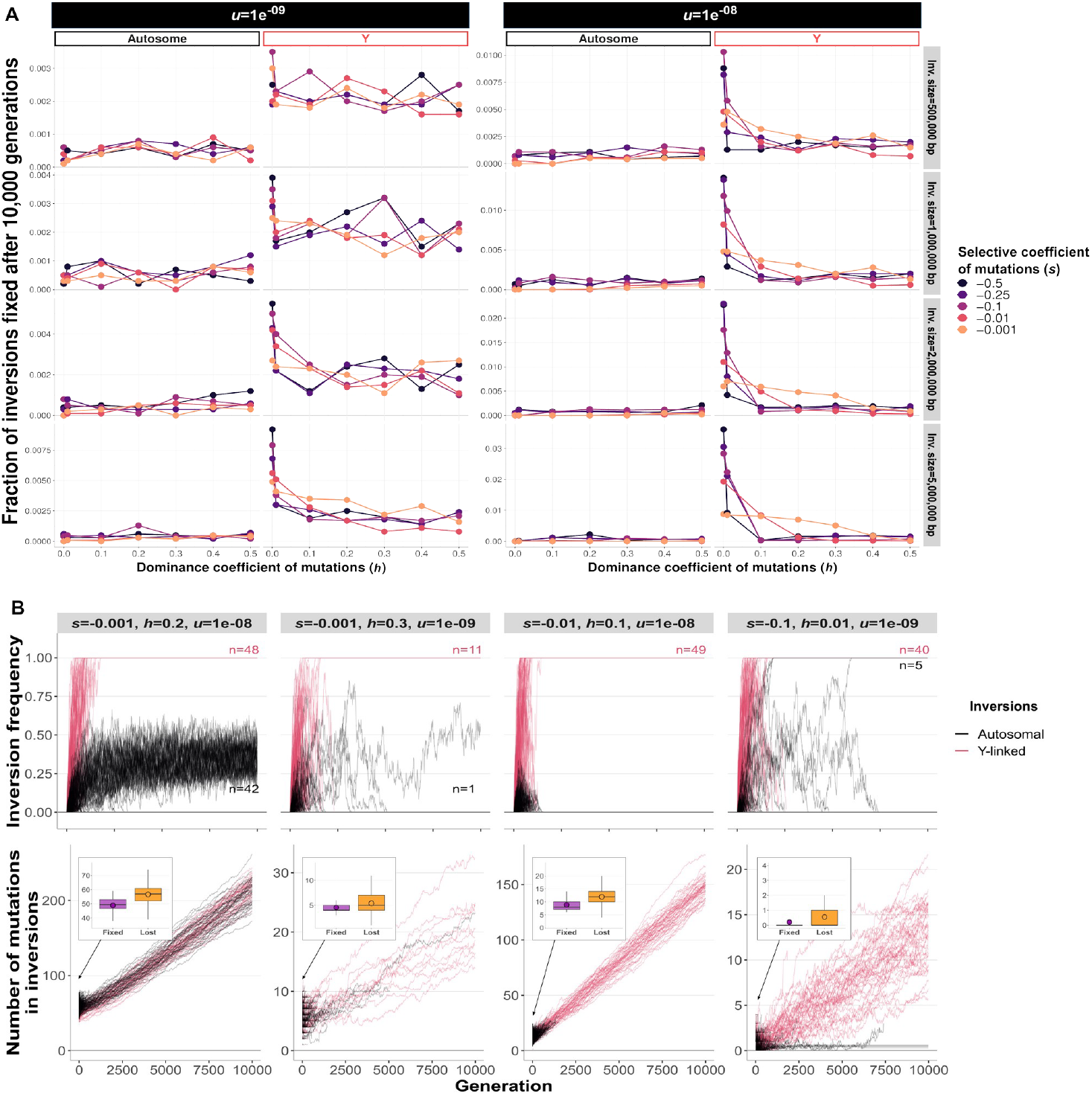
Difference in inversion fates between autosomes and Y-like sex chromosomes due to the sheltering effect. On both autosomes and Y-like sex chromosomes, the spread of inversions is driven by the lower-load advantage (see Jay at al. 2022). However, on Y-like sex chromosomes, the sheltering effect permits inversions to fix despite carrying recessive deleterious mutations, in contrast to autosomes. **A**. Fraction of inversions fixed after 10,000 generations on autosomes and on Y chromosomes depending on the mutation rate (*u*), the selection coefficient of mutations (*s*), the dominance coefficient of mutations (*h*) and the size of the inversions. The figure shows the rates of inversion fixation starting from generation 1. The dataset used is the same as the one used in Jay et al. (2022). A total of 10,000 inversions of 2Mb were studied for each combination of parameter values. Results for inversions of additional inversion sizes are shown in Figure S15 of Jay et al. (2022) and results for additional population sizes are shown in Figure S16 of Jay et al. (2022). Note that the y-axis scales differ from one panel to the other. **B**. Change in inversion frequency and mutation load in stochastic simulations of 1000 individuals under four combinations of parameter values. In these examples, recessive deleterious mutations occur at different rates (*u*=1e-08 or *u*=1e-09), all mutations having the same dominance coefficient (*h*=0.01, 0.1, 0.2 or 0.3) and selection coefficient (*s*=-0.001, -0.01 or -0.1). The data plotted correspond to the same dataset as panel A. For each combination of parameter values, the figures display the frequency and mutation load of 10,000 independent inversions on each of an autosome and a proto-Y chromosome, with each line representing a specific inversion (i.e. a simulation). The evolutionary trajectories of inversions of different sizes, in larger populations, on the X-chromosome or with other parameter combinations are displayed in figures S10-14 in Jay et al. (2022). The inset figures indicate the number of mutations that were present at generation 1 in inversions that were eventually lost and fixed, respectively, showing that fixed inversions initially had fewer mutations than average, i.e. they are those that benefited from an initial lower-load advantage. Boxplot elements: dot: mean, central line: median, box limits: 25th and 75th percentiles, whiskers: 1.5x interquartile range.

While many autosomal inversions with mutation load persisted for hundreds of generations under a wide range of parameters values, the large majority of these autosomal inversions were lost after 10,000 generations (Figure 1B). For instance, with *N*=1000, *s*=0.01 and *h*=0.1, only 73 of out 10,000 inversions of 2 Mb length were still segregating after 500 generations, and they were eventually all lost. These inversions initially spread because they were associated with a lower-than-average mutation load, but their homozygous disadvantage when increasing in frequency prevented them from reaching fixation. Therefore, they remained at intermediate frequencies and they were eventually lost as they accumulated additional deleterious mutations until being more loaded than non-inverted segments (Figure 1B).

In contrast, a substantial proportion of less-loaded inversions capturing the permanently heterozygous sex-determining allele on the Y chromosome spread until they reached fixation within the Y chromosome population, even when not entirely free of deleterious mutations (Figure 1B). For instance, with *s*=0.01, *h*=0.1 and *N*=1000, 49 out of the 10,000 Y-linked inversions of 2 Mb in size became fixed in the Y chromosome population, whereas all autosomal inversions were ultimately lost (Figure 1B). Those 49 inversions increased in frequency because they had a lower load than average (see insets in Figure 1B); then, the sheltering effect, owing to their association to a permanently heterozygous allele, prevented recessive deleterious mutations to impair the fixation of these inversions in the Y chromosome population. Although new deleterious mutations appeared in Y-linked inversions during their spread, they did not accumulate quickly enough to hinder the increase in frequency and fixation of these 49 inversions (Figure 1B).

To check that the higher proportion of inversions fixing on the Y chromosome was not only due to a stronger genetic drift on the Y chromosome, i.e., to the smaller effective population size of these chromosomes relative to autosomes, we performed new simulations and compared the fixation rates of 100,000 inversions on Y chromosomes and autosomes with identical effective population sizes (i.e., autosomes with ¼ of the standard population size; Figure 2). We considered two scenarios: with or without the possibility of fully recessive mutations, both using average dominance coefficients similar to those estimated in nature (0.20–0.30).

**Figure 2.**
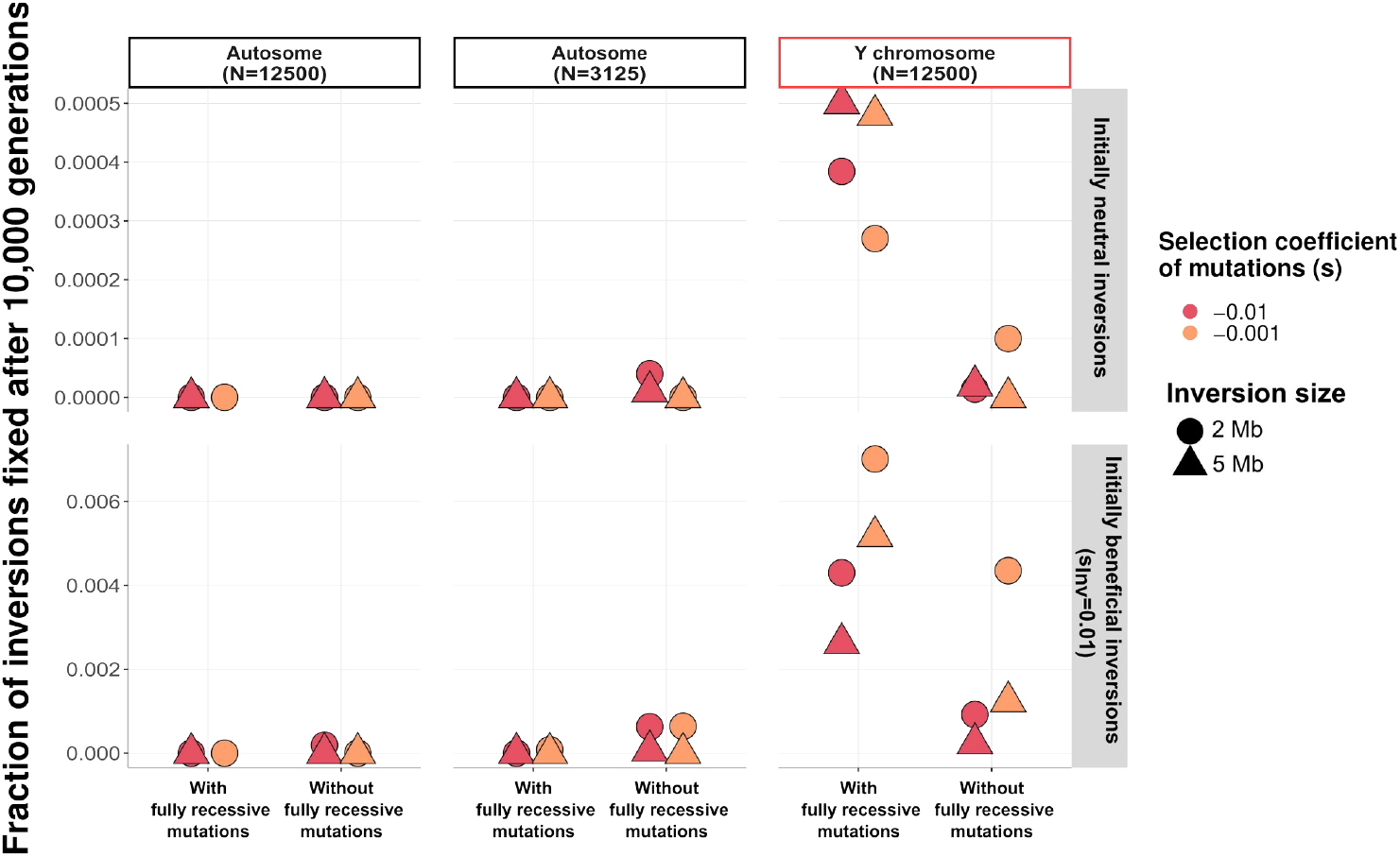
Fraction of inversions fixed after 10,000 generations on autosomes and on Y chromosomes depending on the population size and the intrinsic advantage of inversions. Data from new simulations (i.e. not from Jay et al., 2022). In this second set of simulations, mutations had their fitness coefficients sampled from a gamma distribution of mean -0.001 or -0.01, and their dominance coefficient randomly sampled with uniform probabilities among either [0.1, 0.2, 0.3, 0.4, 0.5] (mean=0.3, no fully recessive mutations), or [0.0, 0.01, 0.1, 0.2, 0.3, 0.4, 0.5] (mean=0.22, with fully recessive mutations). In addition to the simulations of the inversion dynamics on Y chromosomes and on autosomes with identical population sizes (N=12,500), we also performed simulations with an autosomal population of size equivalent to the effective population size of the Y chromosome when total population size is 12,500, i.e. 3,125 (¼ of 12,500). A total of 100,000 inversions of 2Mb and 5Mb were studied for each combination of parameter values. The script used to produce these figures is available at: https://github.com/PaulYannJay/MutationShelteringTheory/tree/main/Reanalyses_bioRxiv. Note that the y-axis scale differs between the different panels.

Our results showed that inversions were more likely to fix on the Y sex chromosome than on autosomes with identical population sizes across 75% of the parameter space explored (Figure 2; note that, due to the limited parameter space explored, this percentage should not be considered as providing an estimate of what occurs in nature). On average, inversions were 35.2 times more likely to fix on Y chromosomes than on autosomes with the same effective population size across the whole range of parameter values studied. As expected, the difference in rates of inversion fixation on autosomes and Y chromosome was stronger when considering mutation landscapes including strongly recessive deleterious mutations, as such scenarios favor the occurrence of the sheltering effect. The population size of autosomes actually had little impact on the probability of inversion fixation: for example, inversions occurring in autosomes were only very slightly more likely to fix (fixation rate of 8e^-6^ *vs* 0.0) when the population size was four times smaller. This is because of the strong disadvantage in fitness when exposing the recessive load at a homozygous state for autosomal inversions. These findings indicate that, while genetic drift plays a small role in the fixation of inversions on Y-like sex chromosomes, the sheltering effect is the primary factor driving the higher fixation rate of inversions on the Y sex chromosome compared to autosomes in our model. This shows that the sheltering effect can be a significant contributor to the evolution of recombination suppression on sex chromosomes.

To illustrate that the sheltering effect and the lower-load advantage are independent effects, we performed similar simulations but with inversions carrying an intrinsic selective advantage other than a lower load (Figure 2). The initial fitness of these inversions depended on both their intrinsic advantage and the random load of deleterious mutations captured at their formation (i.e., a beneficial inversion could here carry a higher-than-average load of deleterious mutations). Without the sheltering effect, we would expect a higher fixation rate of beneficial inversions on autosomes due to more efficient selection generated by the larger effective population size. In contrast, we found that intrinsically beneficial inversions were more likely to fix on the Y chromosomes in 100% of the parameter space, and this was true in both scenarios, with or without fully recessive mutations. Across the parameter values investigated, such inversions were 18.19 times more likely to fix on Y sex chromosomes than on autosomes with an identical population size.

## Discussion

Our analyses of Jay et al. (2022) datasets and of the new simulations provide strong support to our previous conclusion that the combination of the lower-load advantage and the sheltering effect can significantly contribute to the progressive cessation of the recombination on sex chromosome and related genomic architectures: these combined effects promote inversion fixation on Y sex chromosomes, and at higher rates than drift, all other conditions being equal. Our new figures confirmed that including or not the simulations in which the inversions are lost during the first 20 generation has no qualitative effect on our conclusions: inversions remain much more likely to spread and fix on the Y sex chromosome than on autosomes due to the sheltering effect. In fact, this could already be seen in Jay at al. (2022), as the number of inversions lost in the 20 first generations was very similar in autosomes and in the Y sex chromosome (see for example figure S10 in Jay et al. 2022). Therefore, removing the 20 first generations did not introduce any bias for the comparison between autosomes and Y sex chromosomes, which was the goal of our Figure 3C in Jay at al. (2022).

In this manuscript, as well as in Jay et al. (2022), we did not conduct any comparisons with a fully neutral model of sex chromosome evolution, i.e., a model without any deleterious mutations in genomes, where the fixation of inversions would be influenced solely by genetic drift. We consider that such a comparison, which is central in the debate surrounding Jay et al. 2022 study (Olito and Charlesworth 2023), is not relevant. Indeed, we do not think it makes sense to ask whether the sheltering effect and the lower-load advantage leads to higher fixation rates of inversions on sex chromosomes compared to a scenario without any deleterious mutations in genomes. The sheltering effect and the lower-load advantage only arise because the presence of deleterious mutations changes the fitness landscape. The observation that inversions might not be more likely to fix on Y chromosomes for certain parameter values in the presence of deleterious mutations compared to a scenario without any deleterious mutation does not negate the role of the sheltering effect and the lower-load advantage in promoting the fixation of inversions *when deleterious mutations are segregating*, in contrast to previous claims (Olito et al. 2022; Olito and Charlesworth 2023). The sheltering effect is not an intrinsic benefit that can be directly compared to a neutral scenario, as classically done in population genetics. The sheltering effect only mitigates the selective disadvantage incurred by the presence of deleterious mutations.

Thus, s = 0 is not an appropriate benchmark for the processes we analyse in the present paper and in Jay et al. (2022). Deleterious mutations prevent the fixation of most inversions on autosomes (an effect that is absent when s=0), while the sheltering effect protects inversions from this fitness loss on Y-like chromosomes. The sheltering effect thus save inversions specifically on Y-like chromosomes when deleterious mutations are segregating. In contrast, the case s=0 removes both the deleterious effect and the saving effect so that it is not an appropriate control to analyse the sheltering effect (see also Box 3 in Jay at al. 2024 and discussion in Saunders and Muyle 2024). As an analogy, imagine that one wants to test whether one-legged men have a higher probability to cross a mine field without exploding than two-legged men (because we assume that they need twice as fewer steps to cross the field as they jump instead of the step of the missing leg): a comparison of the probabilities to cross the field with and without mines will tell you nothing on the advantage to be one-legged for crossing the field when there are mines. You need to have the mines present (the deleterious effect present) to assess whether being one-legged (being sheltered) gives you an advantage in the real mine field (when deleterious mutations are present). These effects of deleterious mutations are important to take into account as there is ample evidence that genomes harbor numerous deleterious mutations (Eyre-walker & Keightley, 2007).

The appropriate control for genetic drift in our model is an autosome with the same effective population size as the Y sex chromosome, as it retains the effect of deleterious mutations but yields equivalent drift, allowing us to assess whether the higher fixation probability on the Y chromosome is only due to drift. Our new simulations using this correct control show that drift plays a minor role in explaining difference in inversion fixation rate on autosomes and sex chromosomes, the major role being played by the sheltering effect.

### Conditions for the occurrence of the lower-load advantage and the sheltering effect

In this study, we explored a broad parameter space, with *N* ranging from 1,000 to 12,500; *s* from -0.001 to -0.5 (or from 1*Ns* to 500*Ns*); *h* from 0 to 0.5; *u* from 1e-09 to 1e-08; and inversion sizes from 500 kb to 5 Mb. Our results, alongside the analyses in Jay et al. (2022), demonstrate that the number of segregating deleterious mutations and their dominance coefficients are crucial factors for the occurrence of the lower-load advantage and the sheltering effect, with more mutations and more recessive mutations favoring inversion fixation. These quantitative effects are much more informative than an all-or-nothing comparison with no deleterious mutations at all and further show the potential crucial role of deleterious mutations in the evolution of stepwise recombination suppression. Numerous studies in nature have shown that a substantial proportion of new and segregating mutations are deleterious (Eyre-walker & Keightley, 2007). While the precise numbers and effects of these mutations are debated, it is widely accepted that genomes carry tens of thousands of harmful mutations. Therefore, any large inversion (e.g., 1 Mb) is expected to harbor multiple deleterious mutations. For instance, the average human genome contains approximately 4.1 to 5.0 million polymorphic sites, with an estimated 25% of these mutations being deleterious (Racimo and Schraiber, 2014). Even if this estimate was overestimated by a factor of 100, megabase-scale inversions would still contain multiple deleterious variants, setting the stage for the lower-load advantage and the sheltering effect.

Although empirical estimates of the dominance coefficient for mutations in natural populations are limited, studies in *Drosophila*, yeast and nematodes have estimated the average dominance coefficient for deleterious mutations at approximately h = 0.25 (Agrawal et al., 2011; Manna et al., 2011). This suggests that many deleterious mutations have dominance coefficients well below 0.25, in contrast with the assumption of a fixed large h used in analyses by Olito and Charlesworth (2023). For example, Agrawal and Whitlock (2011), using yeast gene knockout data, estimated that mutations affecting catalytic functions have an average dominance coefficient of 0.046. These estimates indicate that the conditions for the sheltering effect are commonly met in natural populations, as we have shown that the effect is more pronounced with mutations having dominance coefficients below 0.2. Similarly, the inversion sizes we modelled are commonly observed in natural populations. For instance, the two most recent evolutionary strata on the human Y chromosome span approximately 1 and 4 Mb, respectively (Zhou et al. 2022).

Finally, it is important to note that evolutionary strata actually rarely evolve in natural populations. For example, the mammalian Y sex chromosome only experienced five successive events of recombination suppression across 180 million years (Cortez et al., 2014, Zhou et al., 2022). Our theory therefore only requires a few lucky inversions that would carry the right combination of mutation number, selective and dominance coefficients to reach fixation, and thus to be able to explain natural patterns of stepwise evolution of recombination suppression.

### Long-term persistence of recombination suppression on sex chromosomes

Another criticism to our previous paper (Jay at al. 2022) was that inversions would still accumulate deleterious mutations after fixation, until a point where selection should favor reversion to a recombining state (Lenormand and Roze 2023). In our previous papers (Jay et al., 2022, 2024), we showed, using simulations and genomic data, that overlapping genomic rearrangements accumulating following recombination suppression could prevent recombination to be restored. Lenormand and Roze (2023) argued that, on the long term and without dosage compensation, the fitness of individuals carrying the inversions having accumulated further load should decrease to a point where species could go extinct, and therefore that the combined effect of the lower-load advantage and sheltering effect could not explain the evolution of recombination suppression on sex chromosomes. However, different types of selection can act at different evolutionary time scales.

For example, it is known for long that selfing or parthenogenesis can be selected for on the short term even if they could be evolutionary dead ends on the long term and lead to species extinction (de Vienne et al., 2013; Wright et al., 2013; Arunkumar et al., 2015). That recombination suppression on sex chromosomes can lead to species extinction unless dosage compensation eventually evolves does not in itself negates the potential role of the sheltering effect in inversion fixation on the short term (see also discussion in Saunders and Muyle 2024). As we have explained in Jay et al. (2024), our theory does not negate a possible role of dosage compensation for the long-term maintenance of sex chromosomes, but it shows that *early* dosage compensation may not be required to explain sex chromosome evolution, in contrast to the conclusions reached in previous studies (Lenormand and Roze, 2022, 2023).

## Conclusion

A reanalysis of the dataset from Jay et al. (2022), along with new simulations, strongly supports our original conclusions and demonstrates that the concerns raised by Olito and Charlesworth (2023) are unfounded. Our conclusions remain robust to the inclusion of inversions lost within the first 20 generations in the analyses and we showed that the comparison with a fully neutral model does not inform on the role of the sheltering effect in contributing to inversion fixation in nature.

Furthermore, we show that, while the lower effective population size of sex chromosomes relative to autosomes may occasionally facilitate the fixation of inversions on Y sex chromosomes, this effect is minimal compared to the sheltering effect in promoting recombination suppression on sex chromosomes across the parameter values studied. As discussed above, we consider that currently available empirical estimates of key parameters -such as population size, inversion size, mutation effect, mutation rate, and mutation dominance-indicate that the combined effects of the lower-load advantage and the sheltering effect are likely to play a substantial role in the evolution of recombination suppression on sex chromosomes and other supergenes in nature. However, substantial uncertainty remains in the estimates and natural variation of these parameters. Future research aimed at refining these estimates could enhance our understanding of the mechanisms driving sex chromosome evolution. In Jay at al. (2024), we also suggested several ways to test the different theoretical models aiming at understanding the evolution of sex chromosomes using experimental or genomic data.

## Acknowledgements

This work was supported by a Human Frontier Science Program (HFSP) fellowship (LT0033/2022-L) to P.J. AV acknowledges support from the chaire program «Mathematical modeling and biodiversity » (Ecole Polytechnique, Museum National d’Histoire Naturelle, Veolia Environnement, Fondation X). We are grateful to Sylvain Billiard and Ariel Offenstadt for their insightful discussions and valuable input.

